# Polypeptides IGF-1C and P24 synergistically promote osteogenic differentiation of bone marrow mesenchymal stem cells in vitro through the p38 and JNK signaling pathways

**DOI:** 10.1101/2021.06.09.447738

**Authors:** Gaoying Ran, Wei Fang, Lifang Zhang, Yuting Peng, Jiatong Li, Xianglong Ding, Shuguang Zeng, Yan He

**Author notes:** Corresponding author, (S. Zeng)., Address: Department of Oral and Maxillofacial Surgery, Stomatological Hospital, Southern Medical University, Guangzhou 510280, China. These authors contributed equally to this work and are co-first authors (Gaoying Ran, Wei Fang).

## Abstract

**Objectives:** Insulin-like growth factor-1 (IGF-1) and bone morphogenetic protein 2 (BMP-2) both promote osteogenesis of bone marrow mesenchymal stem cells (BMSCs). IGF-1C, the C domain peptide of IGF-1, and P24, a BMP-2-derived peptide, both have similar biological activities as their parent growth factors. This study aimed to investigate the effects and their mechanisms of polypeptides IGF-1C and P24 on the osteogenic differentiation of BMSCs.

**Methods:** The optimum concentrations of IGF-IC and P24 were explored. The effects of the two polypeptides on the proliferation and osteogenic differentiation of BMSCs were examined using the Cell Counting Kit-8(CCK-8), Alkaline phosphatase (ALP) staining, ALP activity assay, alizarin red S staining, qPCR, and western blotting. In addition, specific pathway inhibitors were utilized to explore whether p38 and JNK pathways were involved in this process.

**Results:** The optimal concentrations of action were both 50 μg/ml. IGF-1C and P24 synergistically promoted the proliferation of BMSCs, increased ALP activity and the formation of calcified nodules and upregulated the mRNA and protein levels of osterix (Osx), runt-related transcription factor 2 (Runx2), and osteocalcin (Ocn), phosphorylation level of p38 and JNK proteins also improved. Inhibition of the pathways significantly reduced the activation of p38 and JNK, blocked the expression of Runx2 while inhibiting ALP activity and the formation of calcified nodules.

**Conclusions:** These findings suggest IGF-1C and P24 synergistically promote the osteogenesis of BMSCs through activation of p38 and JNK signal pathways.

---

Growth factors have been used in bone tissue engineering and bone defect repair to promote bone formation^[1]^.Insulin-like growth factor-1 (IGF-1) can promote mitosis and is one of the most abundant growth factors that promote the development and metabolism of bone matrix ^[2,3]^.Bone morphogenetic protein 2 (BMP-2) can participate in the differentiation of mesenchymal stem cells (MSCs) into osteoblasts and the formation of bone matrix by osteoblasts^[4]^. These two growth factors have synergistic effects in the process of osteogenic differentiation^[5]^. But the application of growth factors to bone repair is marred by their high cost, their ease of inactivation, and their immunogenicity^[8–10]^. Polypeptides have similar effects as growth factors and can be easily and quickly synthesized on a large scale using a peptide synthesizer. They have high biological activity, good stability, specific function and low immunogenicity, therefore, polypeptides have wide range of applications with a good safety profile^[9,11]^.

IGF-1C, the C domain peptide of IGF-1, and P24, a BMP-2-derived peptide, both have similar biological activities as growth factors^[12,13]^. The C domain of IGF-1, GYGSSSRRAPQT, has been identified as the active region of the IGF-1 protein^[13]^. Feng et al.^[13]^ produced IGF-1C-conjugated chitosan hydrogel and demonstrated that it facilitated the survival of transplanted adipose stem cells. It is still unclear whether IGF-1C can affect the proliferation and differentiation of BMSCs. The P24 peptide (KIPKASSVPTELSAISTLYLSGGC) is truncated from the key amino acid sequence of the BMP-2 “finger epitope”, which can form a binding interface with its receptor to activate the downstream osteogenic signaling pathway^[14]^. In vivo experimental studies have demonstrated that P24-loaded thiolated chitosan/calcium carbonate composite microspheres scaffolds can enhance osteogenesis^[12]^. However, the effect of P24 and IGF-1C on osteogenic differentiation of BMSCs was not studied and the underlying mechanisms of them on osteogenic differentiation were still not clarified.

The mitogen activated protein kinase (MAPK) pathway is the most basic and important signal transduction mediator in various cells; MAPKs mainly include p38 MAPK, c-Jun N-terminal kinase (JNK), and extracellular signal–regulated kinase (ERK). The JNK and p38 signaling pathways play an important role in the regulation of osteogenesis^[6]^. Although Celil et al.^[7]^ suggested that IGF-1 and BMP-2 could act synergistically in MAPK signal transduction, thereby enhancing the expression of osterix (OSX), the P24 and IGF-1C effect of on bone formation via MAPK pathways is still unknown.

Therefore, this study investigated the effects of P24 and IGF-1C on the osteogenic differentiation of BMSCs in vitro. Furthermore, we explored the role of the p38 and JNK signaling pathways in osteogenic differentiation.

## Results

### Optimal concentrations of polypeptides IGF-1C and P24 to promote the proliferation and osteogenic differentiation of BMSCs

The effects of different concentrations of polypeptides on the proliferation and osteogenic differentiation of BMSCs were determined by the CCK-8 and ALP activity assay, respectively. The results showed that there was no cytotoxic effect at peptide concentrations of 0-0.1 mg/ml. Statistical analysis showed that the absorbance values of CCK-8 and ALP activity were significantly higher when the concentrations of these two peptides were both at 50 μg/ml than other concentrations (*P*<0.05) (Figure 1). Therefore, the optimal concentrations of IGF-1C and P24 alone to promote proliferation and osteogenic differentiation of BMSCs were both 50 μg/ml.

**Figure 1.**
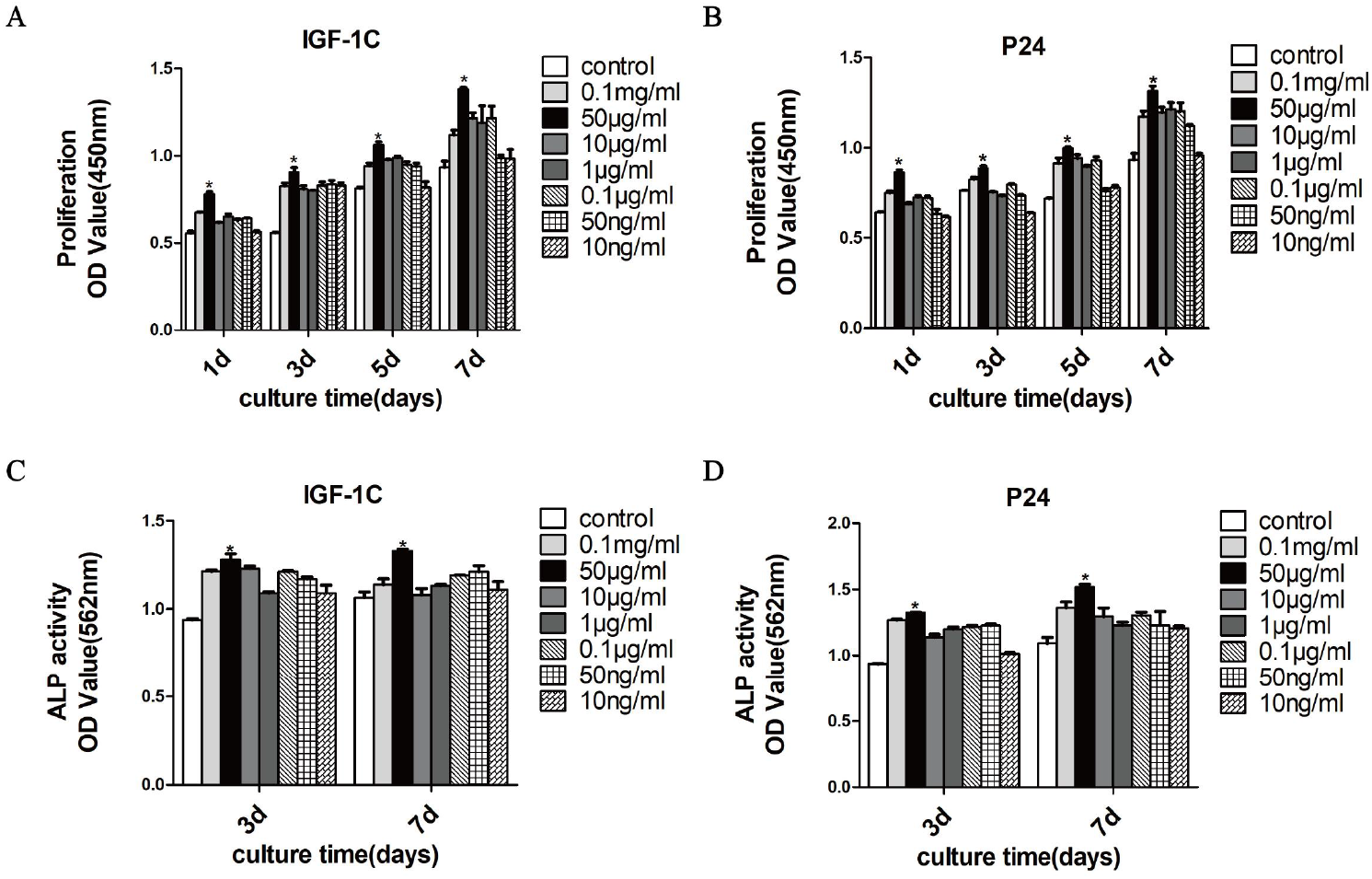
Effects of the concentrations of IGF-1C and P24 on the proliferation and ALP of BMSCs. **A, B** The proliferative activity of BMSCs cultured with different concentrations of IGF-1C or P24 alone for 1, 3, 5, or 7 days. **C and D** The ALP activities of BMSCs cultured with different concentrations of IGF-1C or P24 alone for 3 or 7 days. The proliferation and ALP activity of BMSCs were both highest in the 50 μg/ml groups. The optimal concentration of IGF-1C and P24 was determined to be 50 μg/ml (\**P*<0.05 *vs* other group).

### IGF-1C and P24 synergistically promote the proliferation of BMSCs

After IGF-1C and P24 were cultured alone or in combination with BMSCs for 1, 3, 5, or 7 days, the results showed that, compared with the control group, the IGF-1C and P24 alone groups and the combination group all had more BMSC proliferation. The proliferation ability of the combination group was significantly higher than that of the other groups at a given time *(P* < 0.05) (Figure 2), indicating that both IGF-1C and P24 can promote the proliferation of BMSCs, and these two have synergistic effects when promoting the proliferation of BMSCs.

**Figure 2.**
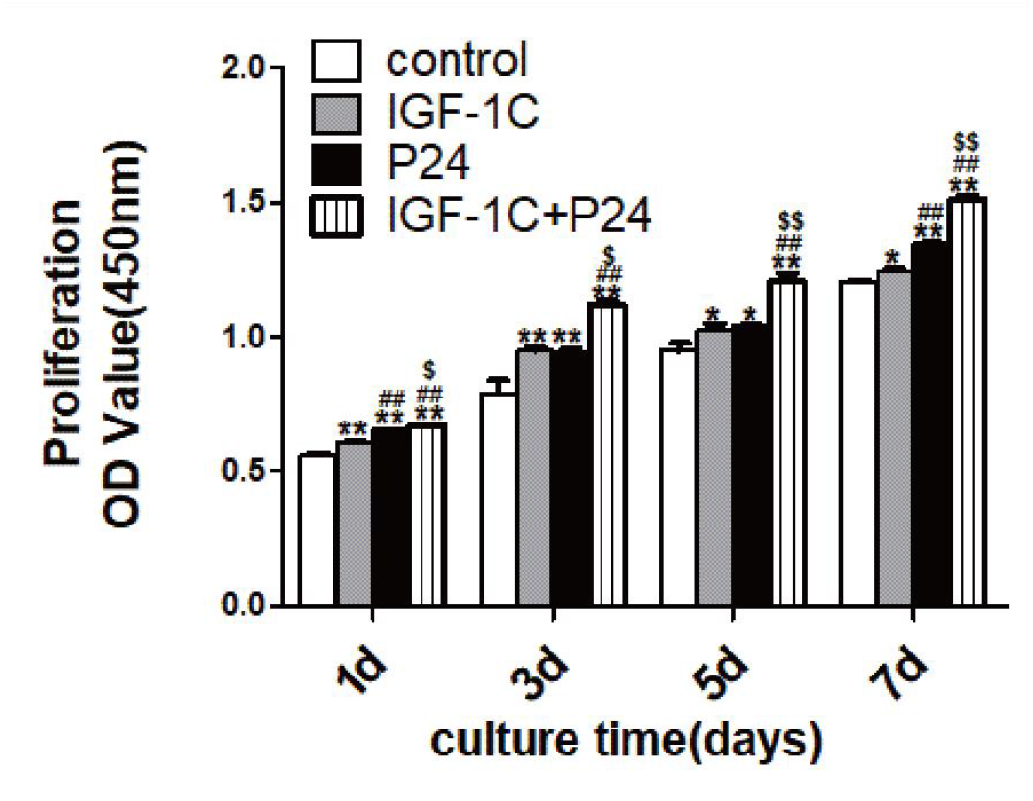
IGF-1C and P24 synergistically promote the proliferation of BMSCs. The results of proliferation activity of BMSCs after 1, 3, 5, or 7 days, when cultured with IGF-1C or P24 alone or their combination showed that the proliferation ability of BMSCs in the combination group was stronger than that of other groups (\**P*<0.05,\*\**P*<0.01 *vs* control; #*P*<0.05,##*P*<0.01 *vs* IGF-1C; $*P*<0.05,$$*P*<0.01 *vs* P24)

### IGF-1C and P24 synergistically promote the osteogenic differentiation of BMSCs

ALP is one of the early markers of osteogenic differentiation of BMSCs. The results of ALP staining showed that dark-bluish-purple-stained substances were formed in all groups on the 3rd and 7th days. However, the ALP-staining BMSCs in the IGF-1C and P24 alone groups and the combination group were significantly higher than those in the control group, and the combination group had significantly more staining than the other groups at each time (Figure 3A-C). The staining results were consistent with the quantitative results of ALP activity (Figure 3D). These results indicate that both IGF-1C and P24 can induce osteogenic differentiation of BMSCs and that these two polypeptides have a synergistic effect in promoting the osteogenic differentiation of BMSCs.

**Figure 3.**
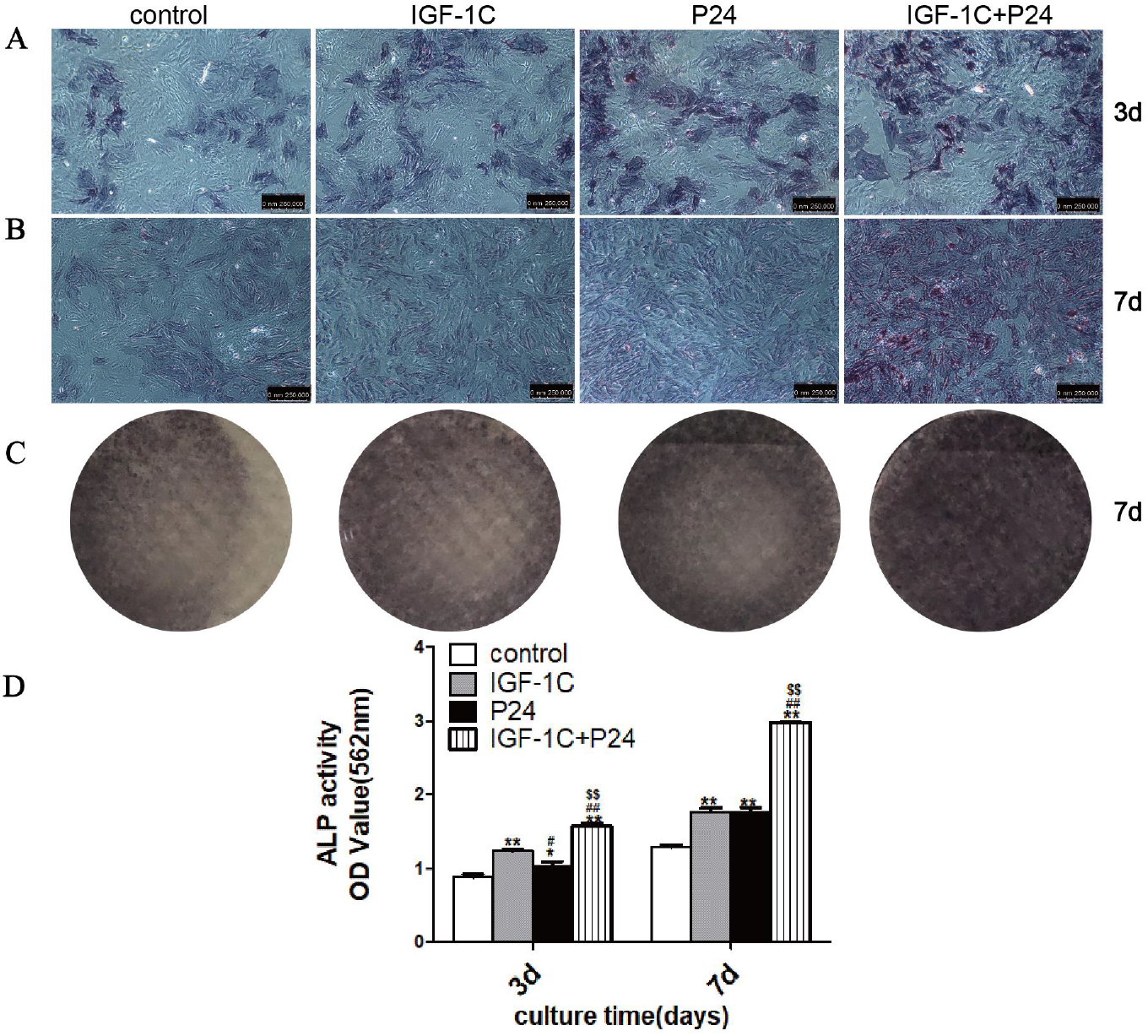
IGF-1C and P24 synergistically promote the osteogenic differentiation of BMSCs. **A, B** The ALP staining images of culture with IGF-1C or P24 alone or their combination for 3 days and 7 days under a microscope (×100). **C** Gross images of ALP staining of cultures with IGF-1C or P24 alone or their combination for 7 days. **D** Quantitative detection of ALP activity after culture with IGF-1C or P24 alone or their combination for 3 days and 7 days. The results showed that culture with IGF-1C or P24 alone or their combination increased the ALP activity of BMSCs, and the combination of IGF-1C and P24 was stronger than the other three groups (\**P*<0.05,\*\**P*<0.01 *vs* control; #*P*<0.05,##*P*<0.01 *vs* IGF-1C; $*P*<0.05,$$*P*<0.01 *vs* P24).

### IGF-1C and P24 synergistically promote the mineralization of BMSCs

Calcified nodules are one of the markers of the osteogenic differentiation of BMSCs at the late stage. ARS staining results showed that red calcified nodules were formed in all groups when cultured with IGF-1C or P24 alone or both together for 14 or 21 days. The number and area of calcified nodules in BMSCs of the IGF-1C and P24 alone groups and the combination group were higher than those in the control group (Figure 4A-C), and the combination group had significantly more than the alone groups at each time. The staining results were consistent with the semiquantitative results of ARS (Figure 4D), indicating that both IGF-1C and P24 induced mineralization in BMSCs, and the two polypeptides had synergistic effects on the mineralization induction of BMSCs

**Figure 4.**
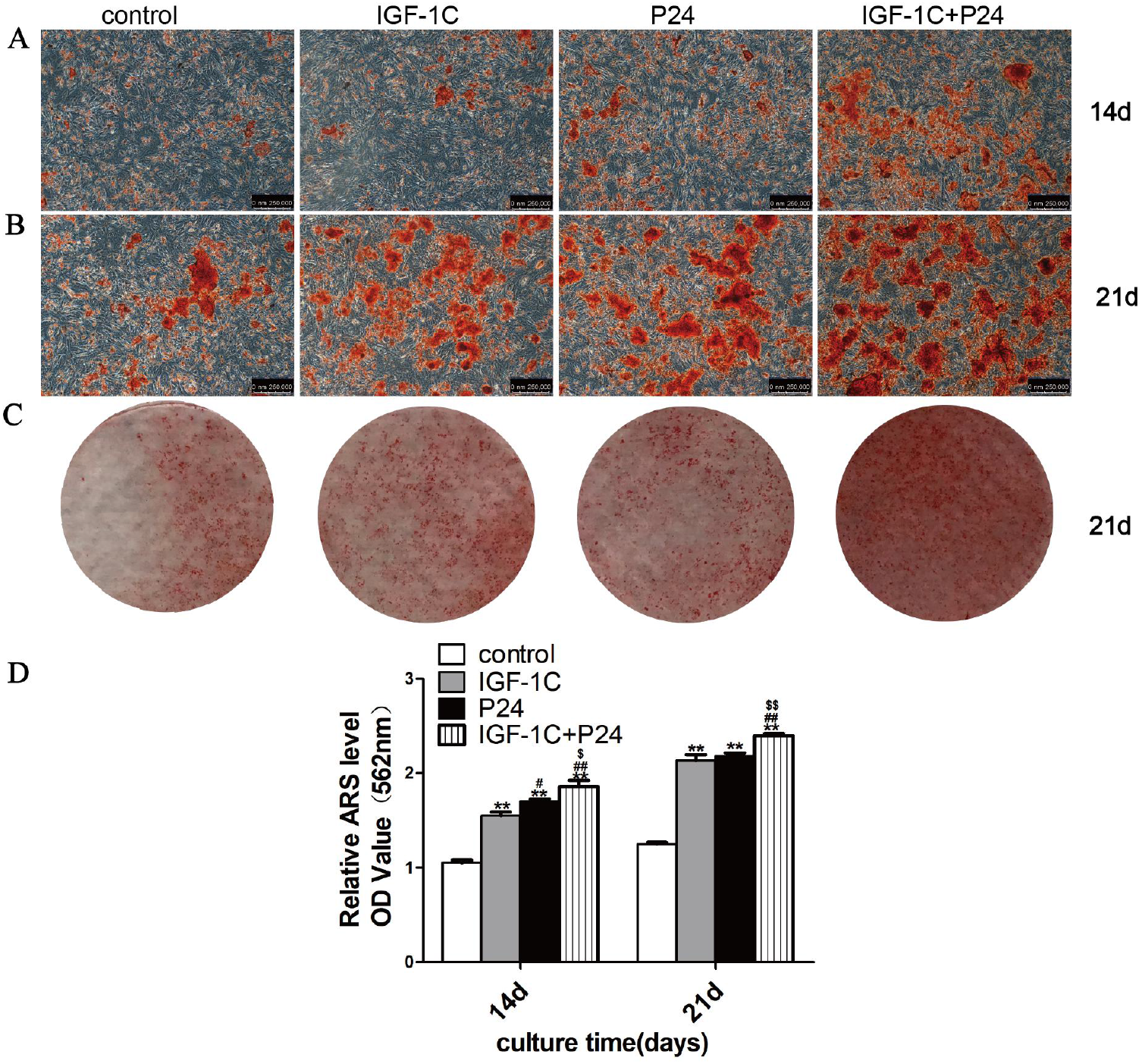
IGF-1C and P24 synergistically promote the mineralization of BMSCs. **A, B** The ARS staining images of BMSCs cultured with IGF-1C or P24 alone or both together after 14 or 21 days under a microscope (×100). **C** Gross image of ARS staining in BMSCs after culture with IGF-1C or P24 alone or their combination for 21 days. **D** Semiquantitative detection of ARS staining of culture with IGF-1C or P24 alone or their combination in BMSCs for 14 or 21 days. The results showed that both IGF-1C and P24 alone and in combination promoted the formation of mineralized nodules in BMSCs, and the mineralization effect of the combination group of IGF-1C and P24 was stronger than that of the other three groups. The semiquantitative results of ARS were consistent with those of the staining results (\**P*<0.05,\*\**P*<0.01 *vs* control; #*P*<0.05,##*P*<0.01 *vs* IGF-1C; $*P*<0.05,$$*P*<0.01 *vs* P24).

### IGF-1C and P24 synergistically up-regulated osteogenesis-related gnens in BMSCs

qPCR detected the gene expression of Osx, Runx2 and Ocn. The results showed that after 7, 14, and 21 days of culture (Figure 5), the mRNA levels of Osx, Runx2 and Ocn were all higher in the IGF-1C and P24 alone and the combination groups than the control group. The expression level were the highest in the combination group, indicating that both IGF-1C and P24 could induce bone formation in BMSCs, and these two polypeptides had a significant synergistic effect in inducing bone formation.

**Figure 5.**
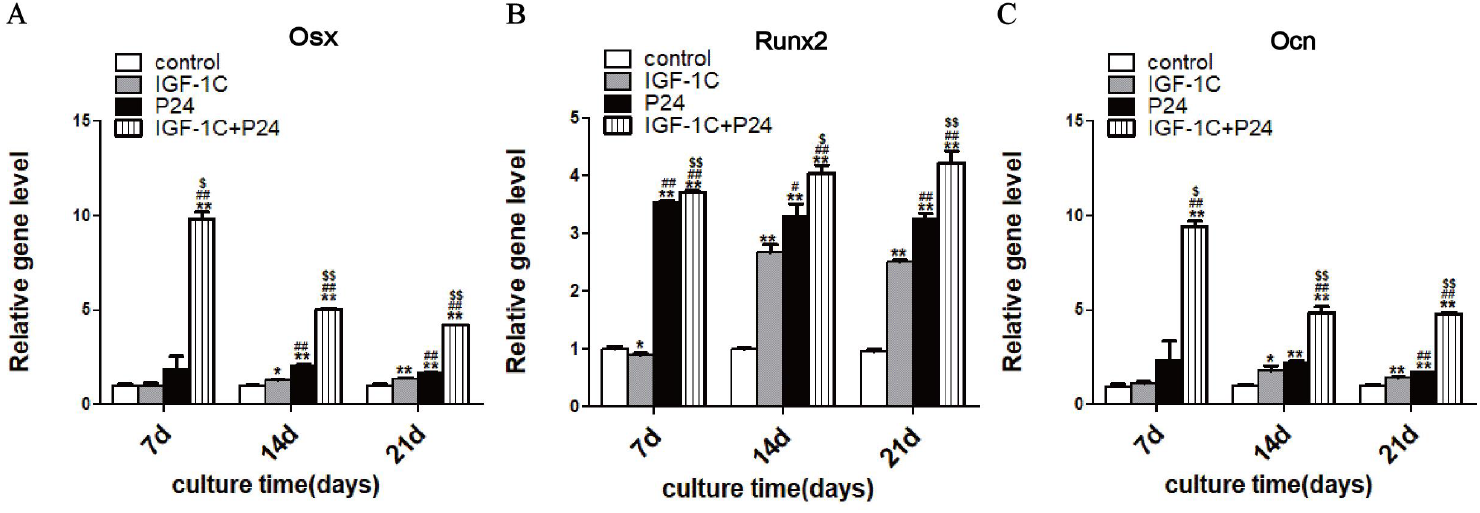
IGF-1C and P24 synergistically up-regulated osteogenesis-related gnens in BMSCs. **A, B, and C** The results of Osx,Runx2, and Ocn gene expression after culture with IGF-1C or P24 alone or their combination for 7, 14, or 21 days showed that the expression of the Osx,Runx2, and Ocn genes was significantly increased when IGF-1C and P24 were cultured in combination compared with other treatments (\**P*<0.05,\*\**P*<0.01 *vs* control; #*P*<0.05,##*P*<0.01 *vs* IGF-1C; $*P*<0.05,$$*P*<0.01 *vs* P24).

### IGF-1C and P24 synergistically up-regulated osteogenesis-related proteins in BMSCs

Western blot was used to detect the expression levels of the osteogenesis-related proteins Runx2, OCN and OSX. The results showed that the expression levels of Runx2, OCN, and OSX in BMSCs of the IGF-1C and P24 alone and the combination groups all increased after 7, 14, and 21 days compared with the control group. In addition, the expression level of the combination group was significantly higher than that of the other three groups, which also indicated that IGF-1C and P24 had a significant synergistic effect in inducing osteogenesis. (Figure 6).

**Figure 6.**
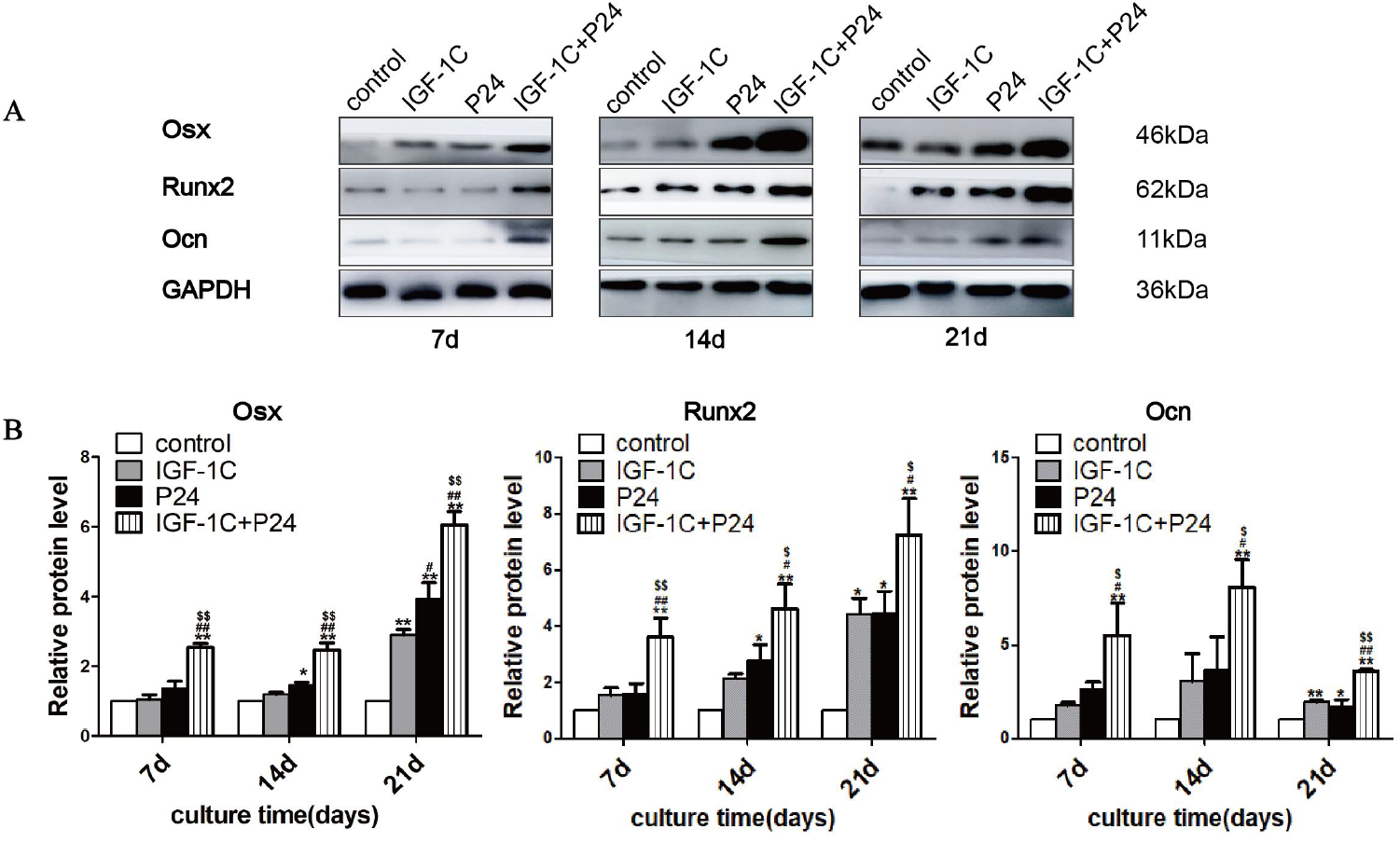
IGF-1C and P24 synergistically up-regulated osteogenesis-related proteins in BMSCs. **A** Western blot detection of the expression levels of osteoblast differentiation-associated proteins after culture with IGF-1C and P24 alone and in combination for 7, 14, and 21 days. **B** Quantitative analysis of Osx, Runx2 and Ocn protein levels; The results showed that when the protein levels of the internal reference GAPDH were essentially the same, combination culture of IGF-1C and P24 caused a significant increase in the expression levels of Osx, Runx2 and Ocn proteins compared to the other three groups (\**P*<0.05,\*\**P*<0.01 *vs* control; #*P*<0.05,##*P*<0.01 *vs* IGF-1C; $*P*<0.05,$$*P*<0.01 *vs* P24)

### IGF-1C and P24 synergistically enhanced phosphorylation of p38 and JNK in BMSCs

The phosphorylation of p38 and JNK plays an important role in the regulation of osteogenic differentiation of BMSCs. Western blot was used to detect the expression of phosphorylated p38 (p-p38) and phosphorylated JNK (p-JNK) in BMSCs. After 2, 4, and 6h of stimulation, both IGF-1C and P24 activated p-p38 and p-JNK in BMSCs compared with their absence, and both p-p38 and p-JNK protein levels were higher in the combined group than the other groups (Figure 7). These results indicate that both IGF-1C and P24 can activate the p38 and JNK signaling pathways during the process of osteogenic differentiation, and these two polypeptides have a synergistic effect in activating the p38 and JNK signaling pathways.

**Figure 7.**
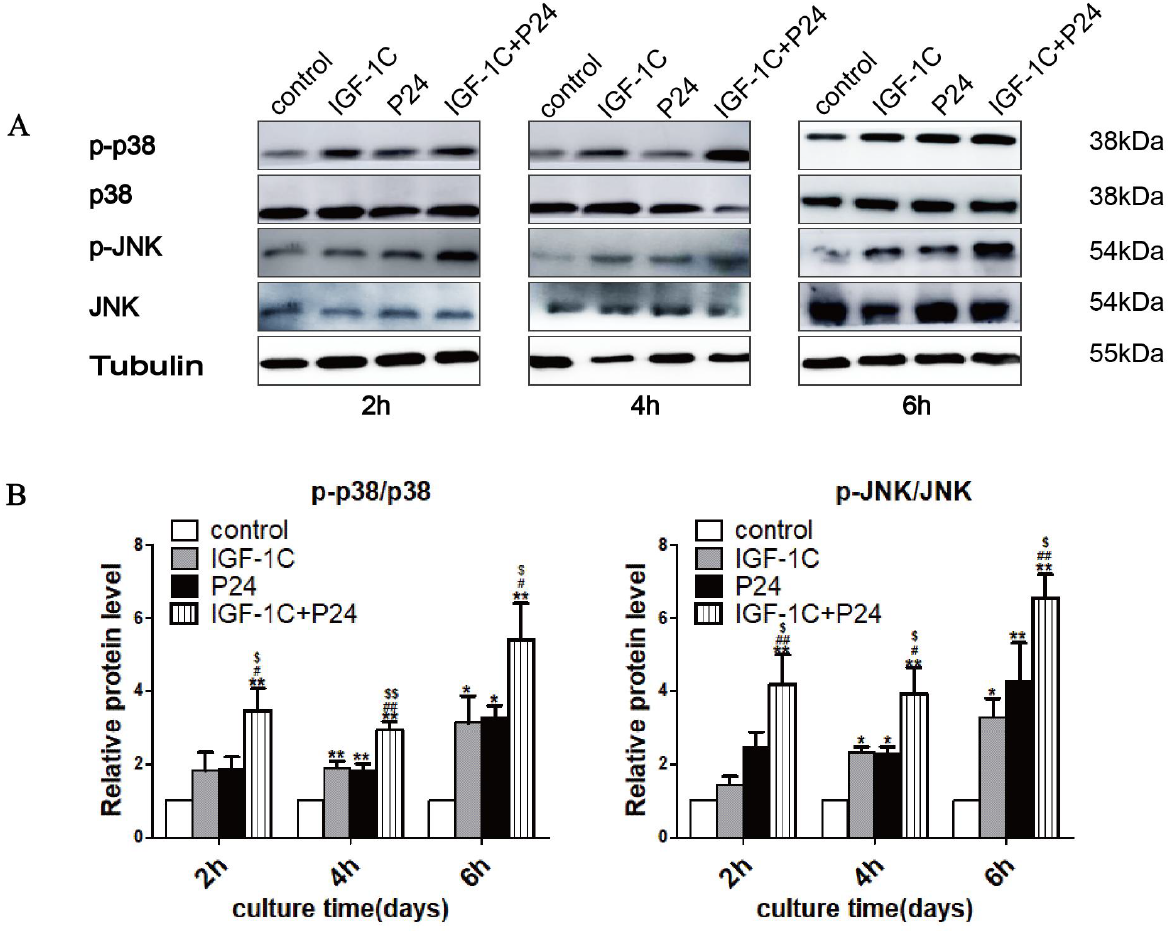
IGF-1C and P24 synergistically enhanced phosphorylation of p38 and JNK in BMSCs. **A** Western blot detection of JNK and p38 after culture with IGF-1C or P24 alone or both together for 2, 4, or 6 h. **B** Quantitative analysis of the expression levels of p38 and JNK phosphorylation. The results showed that the total protein levels of p38 and JNK and the protein level of the internal control tubulin were basically the same after 2, 4, and 6 h of culture, while the expression levels of p-38 and p-JNK were higher after 2, 4, and 6 h of combination culture compared with the other cultures. (\**P*<0.05,\*\**P*<0.01 *vs* control; #*P*<0.05,##*P*<0.01 *vs* IGF-1C; $*P*<0.05,$$*P*<0.01 *vs* P24).

### The p38 and JNK inhibitors suppressed polypeptides – mediated osteogenic differentiation enhancement of in BMSCs

SB203580 and SP600125 are specific inhibitors of the p38 and JNK signaling pathways, respectively. At 4 h, compared with the control group and the IGF-1C+P24 combination group, the SB203580 and SP600125 groups had significantly less p-p38 and p-JNK, and the expression of the Runx2 was downregulated. These results were consistent with the results of ALP and ARS staining (Figure 8). They preliminarily indicated that both IGF-1C and P24 could regulate the osteogenic differentiation of BMSCs through the activation of the p38 and JNK signaling pathways and that IGF-1C and P24 have synergistic effects in activating these two pathways.

**Figure 8.**
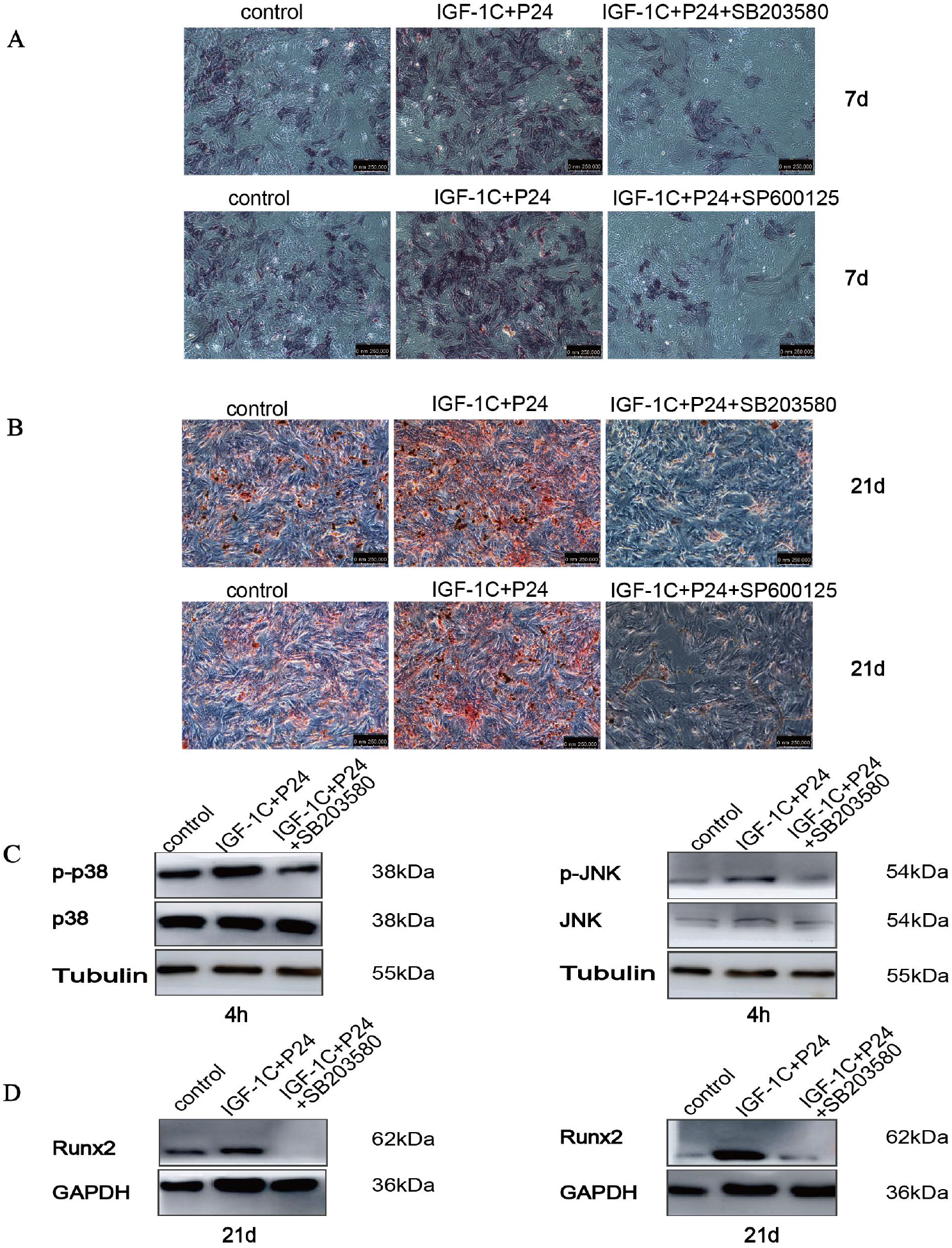
The p38 and JNK inhibitors suppressed polypeptides-mediated osteogenic differentiation enhancement of in BMSCs. **A** ALP staining image of combination culture with IGF-1C and P24 with or without the addition of SB203580 or SP600125 for 7 days (×100). **B** ARS staining image of combination culture with IGF-1C and P24 with or without the addition of SB203580 or SP600125 for 21 days (×100). **C** The expression levels of p38 and JNK in combination culture with IGF-1C and P24 with or without the addition of SB203580 or SP600125 for 4 h. **D** The expression level of the osteoblast differentiation-associated protein Runx2 in combination culture with IGF-1C and P24 with or without the addition of SB203580 or SP600125 for 21 days. SB203580 or SP600125 inhibited the effect of IGF-1C and P24 combination culture on the levels of p-p38, p-JNK, and Runx2. The ALP and ARS staining also showed that the addition of SB203580 or SP600125 inhibited the activity of ALP and the formation of calcified nodules.

## Discussion

Under physiological conditions, bone repair and reconstruction are extremely slow and complex processes. Inducing BMSCs to differentiate into bone cells may be an effective strategy to treat bone defects and promote fracture healing^[15]^. Osteogenic differentiation involves many types of intercellular and intracellular signal transduction, such as signaling pathways, transcription factors, and growth factors, which coordinately regulate bone metabolism^[4,16]^. Many cytokines play important roles in the process of osteogenesis, including transforming growth factor-β (TGF-β), BMPs, platelet-derived growth factor, and IGF^[1,17,18]^. The regulatory function of local cell growth factors during the osteogenesis process amounts to a network regulation of growth factors. As research on bone tissue engineering technology has advanced, one research hotspot that has emerged is how to combine the application of growth factors with different bioactivities to promote bone regeneration in order to mimic the natural osteogenesis process and improve osteogenic efficiency. BMP-2 has a strong osteogenic induction ability, and IGF-1 can also promote the proliferation and differentiation of BMSCs, so both can promote bone formation. IGF-1C, the C domain peptide of IGF-1, and P24, a BMP-2-derived peptide, both have similar biological activities as growth factors. However, it is unclear whether these two active polypeptides can promote the proliferation and differentiation of BMSCs in vitro. This study aimed to investigate whether there is a synergistic effect of IGF-1C and P24 in promoting osteogenesis and to preliminarily explore the underlying mechanism.

The ability of a growth factor to promote stem cell proliferation and differentiation depends heavily on its concentration^[8]^. Likewise, the optimal concentration of polypeptide promoted its biological activity; In contrast, high concentrations inhibited the proliferation of stem cells^[19,12]^. The optimal concentrations of IGF-1C and P24 for promoting BMSC proliferation in vitro were still unclear.

We first examined the effects of the concentrations of these two polypeptides on the proliferation and differentiation of BMSCs. The results of CCK-8 and ALP activity assays showed that when IGF-1C and P24 were used alone, they had no cytotoxicity. They had similar activities as growth factors and could promote the proliferation and differentiation of BMSCs. This ability to promote proliferation and differentiation was related to their concentrations. Within the concentration range used here, 50 μg/mL of both polypeptides maximally promoted the proliferation and differentiation of BMSCs, so 50 μg/mL was used in all the experiments.

Compared either one alone, BMP-2 and IGF-1 together can enhance wound healing and tissue regeneration synergistically^[5]^. IGF-1 stimulates the growth and proliferation of osteoblasts and enhances local bone integration. It can also induce ALP activity, upregulate Ocn expression, and increase matrix calcium content and nodule formation in osteoblasts^[2,3]^. IGF-1 is involved in osteoblast maturation and mineralization in the process of bone repair^[20, 21]^. The growth factor BMP-2, which plays an important role in osteogenic differentiation and fracture repair, belongs to the TGF-β superfamily^[11]^. The addition of BMP-2 to the culture medium can rapidly induce the proliferation of mouse skeletal stem cells^[22]^. Cheng et al.^[23]^ found that BMP-2 is one of the most effective factors at inducing BMSCs to differentiate into osteoblasts, and it can stimulate ALP and matrix calcification^[24]^. Chen et al.^[25]^ reported that the combined administration of BMP-2 and IGF-1 resulted in the highest ALP activity in periodontal ligament fibroblasts. Kim et al.^[5]^ prepared a chitosan gel/gelatin microsphere dual delivery system to control the release of BMP-2 and IGF-1; combined delivery has significantly enhanced ALP activity, wound healing, and tissue regeneration in early osteoblasts compared to single growth factor release. Therefore, we tested IGF-1C and P24 alone and in combination were used to culture with BMSCs in vitro. According to the statistical analysis of the CCK-8 assay results, after 1, 3, 5, and 7 days of culture, compared with their absence, both IGF-1C and P24 alone promoted the proliferation of BMSCs, but the combination of the two synergistically promoted the proliferative activity of BMSCs.

In the process of repairing bone defects, stem cells need to undergo a process of directed migration and osteogenic differentiation. Under in vitro culture conditions, the formation of calcified bone-like tissues by BMSCs includes three stages: the cell proliferation phase, cell aggregation and secretion phase, and the extracellular matrix maturation and mineralization phase. In the first stage, stem cells differentiate into osteoblasts and express Runx2^[26]^. Runx2 is a core transcription factor that can induce the transformation of immature osteocytes to mature osteocytes at early stage^[27]^. In this study, the mRNA and protein levels of Runx2 were upregulated under combination culture with IGF-1C and P24, more so than under IGF-1C or P24 culture alone. In the second stage, during the secretory phase, the level and activity of ALP increased to promote cell differentiation and extracellular matrix maturation^[18]^. In this study, ALP was upregulated when cultured with IGF-1C or P24 alone, but ALP activity was the highest with combination culture. In the third stage, during the mineralization phase, osteoblasts express Ocn to promote the formation of calcium nodules. Ocn is the most abundant noncollagenous bone matrix protein and is only expressed in the late stage of osteogenic differentiation. Therefore, the expression of Ocn in osteoblasts is used as a marker of osteoblast metabolic activity and mineral deposition^[28]^. Combined with ARS staining and the expression of Ocn genes and proteins, both IGF-1C and P24 synergistically promoted the osteogenic differentiation of BMSCs into the late stage. A downstream target of Runx2, Osx is necessary for bone formation and osteoblast differentiation. It can participate in both early osteogenic differentiation and late osteogenesis. The expression of OSX protein and mRNA was consistent with the expression of early Runx2. It was also consistent with the expression of late Ocn. In the entire differentiation process, these factors acted synergistically to ensure the normal maturation of the extracellular matrix, which indicates that IGF-1C and P24 synergistically promote the osteogenesis of BMSCs, and the synergistic osteogenic effect of these two peptides occurs throughout all stages of osteogenesis.

The MAPK signaling pathway is one of the most important signal transduction systems and is widely involved in cell growth, proliferation and differentiation^[9]^. During bone development, bone formation, and bone homeostasis, p38 can be activated by BMP-responsive kinase to regulate the differentiation of MSCs^[30, 31]^. JNK signaling can promote cell growth and differentiation and can transduce extracellular signals to nuclear transcription factors to enhance transcription and promote osteogenesis^[29,32]^. JNK and p38 are commonly referred to as stress-activated protein kinases, which together promote the synthesis of extracellular matrix and the deposition of calcium^[29]^. IGF-1 can trigger MAPK signaling pathway^[7,32]^, BMP-2 can also act through a nonclassical pathway that involves MAPK signaling pathways to directly regulate the expression of Osx, a key transcription factor in osteogenesis^[33]^. Experiments had shown that IGF-1 can upregulate BMP-2 expression, and the upregulation of BMP-2 expression by IGF-1 is achieved through MAPK signaling^[34]^. MAPK signaling was also found to be the hub that mediates the induction of osteogenesis in MSCs by BMP-2 and IGF-1^[7,33]^. Therefore, we investigated the role of the p38 and JNK pathways in the regulation of IGF-1C- and P24-mediated osteogenic differentiation of BMSCs. The results showed that both IGF-1C and P24 increased p-p38 and p-JNK in BMSCs, which were the highest when the two polypeptides were combined. Here, SB203580 and SP600125 inhibited the activity of ALP and the formation of mineralized nodules while downregulating the protein level of Runx2 by inhibiting the activation of the p38 and JNK pathways.

Taken together, IGF-1C and P24 can synergistically activate the p38 and JNK signaling pathways in BMSCs, thereby increasing ALP activity; upregulating Runx2, Ocn, and Osx gene and protein expression; and promoting the formation of calcified nodules.

## Conclusion

In conclusion, IGF-1C and P24 can activate the p38 and JNK signaling pathways in BMSCs, which means that both pathways play a crucial role in the synergistic induction of osteogenesis in BMSCs by IGF-1C and P24. This study can provide a foundation for tissue engineering–based repair of bone defects.

## Experimental procedures

### Grouping

1. BMSCs were cultured in peptide-free osteogenic induction medium as controls. The concentrations of polypeptides IGF-1C/P24 were divided into eight groups: control, 10 ng/ml, 50 ng/ml, 0.1 μg/ml, 1 μg/ml, and 10 μg/, 50 μg/ml, and 0.1 mg/ml. The optimal concentrations of IGF-1C and P24 for promoting proliferation and osteogenic differentiation when they were added individually or jointly to BMSCs culture were determined using the CCK-8 and ALP activity assays.
2. After the optimal concentrations of polypeptides IGF-1C and P24 were determined, BMSCs cultured with peptide-free osteogenic induction medium served as controls. Cells were divided into the control group, the IGF-1C group, the P24 group, and the IGF-1C+P24 group. The effects of polypeptides IGF-1C and P24 alone and in combination on the proliferation and osteogenic differentiation of BMSCs were determined using CCK-8, ALP activity detection, ARS staining, Western blot, and qPCR assays.
3. After the addition of the p38 inhibitor SB203580 (25 μM, Selleck, S1076, USA) and the JNK inhibitor SP600125 (10 μM, Selleck, S1460, USA), BMSCs were cultured in osteogenic induction medium. Cells without peptides and inhibitors and served as controls. Cells were divided into the control group, the combination group, and the inhibitor+IGF-1C+P24 group. The mechanism of IGF-1C and P24 in promoting osteogenic differentiation of BMSCs was preliminarily investigated using ALP staining, ARS staining, and Western blotting.

### Materials

Second-generation Sprague-Dawley rBMSCs were purchased from the Cell Bank of the Chinese Academy of Sciences (catalogue No. SCSP-402) and were cultured according to the instructions. BMSCs have been verified by the Chinese Academy of Sciences for their adipogenic, cartilaginous, and osteogenic potential. P24 and IGF-IC were purchased from Shanghai ZiYu Biotech Co., Ltd., China. BMSC osteogenic induction medium (Cyagen Biosciences Inc., RASMX-90021, China) was used.

### Methods

#### Culture of BMSCs

BMSCs in F12 (Gibco, 31331093, US) containing 10% FBS (Gibco, 10270-106, US) and 1% penicillin-streptomycin (Gibco, 15140-122, US) were cultured in a 37°C, 5% CO_2_ humidified environment. When BMSCs reached more than 80% confluence, they were switched to the osteogenic induction media described in section grouping. The medium was changed once every 3 days.

#### CCK-8 assay

The effect of polypeptides on the proliferation of BMSCs was detected using CCK-8. After BMSCs were cultured for the specified time, F12 containing 10% CCK-8 working solution (Beyotime, C0042, China) was added and incubated at 37°C for 1 h. The absorbance of each well at 450 nm was recorded using a microplate reader.

#### ALP activity assay

The ALP assay was used to quantify the ALP activity of BMSCs. The cells were washed with PBS and fixed with 70% ethanol for 10 min. The ALP staining kit (Beyotime, C3206, China) was used for staining. Then,the ALP activity was assessed by the ALP assay kit (Beyotime, P0321M, China). The protein concentration was determined using BCA protein assay reagent (Beyotime, P0012, China) and the absorbance was measured at 562 nm.

#### ARS staining

ARS staining was used to detect the formation of calcified nodules in BMSCs. At the specified time, the cells were rinsed with PBS, fixed in prechilled 70% ethanol for 15 min, and stained with ARS at room temperature (Cyagen Biosciences Inc., PH-140, China) for 30 min. For quantitative analysis, they were eluted with 10% cetylpyridinium chloride and stained with ARS extracted from cell matrix. The absorbance was measured at 562 nm.

#### qPCR analysis

At the specified time, total RNA was extracted using TRIZOL reagent (Invitrogen, 15596-018, USA). Total RNA was reverse-transcribed using the PrimeScript™RT kit (Takara, RR047A, Japan) to obtain cDNA. cDNA products were amplified by qPCR using the SYBR^®^Premix Ex *Taq* TM kit (Takara, RR420A, Japan) in an Applied Biosystem 7500 Fast Real-Time PCR System under standard PCR conditions. Glyceraldehyde-3-phosphate dehydrogenase (GAPDH) was the internal control. The mRNA transcription level of each target gene was quantified by the 2^-ΔΔCt^ method.The primers for Osx, Ocn, Runx2 and GAPDH were as follows: Osx (F: 5’-CTGCTTGAGGAAGAAGCTC – 3’ and R: 5’ – TATGGCTTCTTTGTGCCTC – 3’); Runx2 (F: 5’ – GAACCAAGAAGGCACAGAC – 3’ and R: 5’ – AATGCGCCCTAAATCACTG-3’); Ocn (F: 5’-TATGGCACCACCGTTTAGGG-3’ and R: 5’ – CTGTGCCGTCCATACTTTCG – 3’); GAPDH (F: 5’ – GGCAAGGTCATCCCAGAGCT-3’ and R: 5’-CCCAGGATGCCCTTTAGTGG-3’).

#### Western blot

Western blot assays were run to detect the expression of osteogenic-related proteins and MAPK signaling pathway proteins in BMSCs. BMSCs were cultured for the specified time and then lysed in RIPA^TM^ lysis buffer. And then the protein concentration was determined using the BCA reagent kit. The protein extracts were subjected to SDS-PAGE electrophoresis, transferred to a 0.22-μm nitrocellulose membrane, and blocked with milk for 2 h. The membranes were respectively incubated overnight at 4°C with anti-Osx (1: 800: AffinityDF7731China), anti-Runx2 (1:1000, AF5186, DF7731, China), anti-Ocn (1:800, Affinity, DF12303, China), anti-p38(1:1000, CST, 9212, US), anti-p-p38 (1:1000, CST, 4511, US), anti-Jnk (1:1000, CST, 4668, US), anti-p-JNK (1:1000, CST, 9351, US), anti-GAPDH (1:1000, Abcam, ab8245, US), and anti-Tubulin (1:1000, Abcam, ab7291, US). Then, were incubated with secondary antibody horseradish peroxidase (1:5000, Boster Biological Technology Co. Ltd, BA1054, China) for 2 h. Finally, an enhanced ECL reagent kit (CST, 6883S, US) and GE ImageQuant LAS 500 exposure system were used for detection, and ImageJ software was used for quantitative analysis.

#### Statistical analysis

The results were representative of at least three independent experiments, and the data are expressed as the mean ± standard deviation. SPSS 21.0 software was used to perform analysis of variance on data among multiple groups. *P* < 0.05 was considered significant.

## Acknowledgements

The work was financially supported by Guangzhou Science and Technology Project of China (201707010193)and the Intra-hospital Fund of Southern Medical University of Stomatological Hospital (PY2020021)..

## Author Contributions

Conceived and designed, Shuguang Zeng; performed the experiments, Gaoying Ran and Wei Fang; analyzed the data, Gaoying Ran,Lifang Zhang and Yuting Peng; wrote the paper,Gaoying Ran,Wei Fang and Shuguang Zeng; All authors discussed the results and implications and commented on the manuscript at all stages. All authors have read and agreed to the published version of the manuscript.

## Data availability statement

Raw data were generated at Laboratory of Heart Center, Guangdong Provincial Biomedical Engineering Technology Research Center for Cardiovascular Disease, Zhujiang Hospital, Southern Medical University. Derived data supporting the findings of this study are available from the corresponding author, Shuangguang Zeng, on request.

